# APPENDICULAR MORPHOLOGY AND LOCOMOTOR PERFORMANCE OF TWO MORPHOTYPES OF CONTINENTAL ANOLES: *Anolis heterodermus* AND *Anolis tolimensis*

**DOI:** 10.1101/743807

**Authors:** Juan Camilo Ríos-Orjuela, Juan Sebastián Camacho-Bastidas, Adriana Jerez

## Abstract

*Anolis* lizards have been a model of study in ecomorphology in the Caribbean islands because species with the same type of microhabitat share similar morphological features. But despite their great diversity, little is known about continental species. We analyzed the relationship between the anatomical characteristics of the appendicular skeleton and the locomotor performance of two *Anolis* species found in Colombia that have different use of habitat. *Anolis heterodermus* a strictly arboreal species was compared with *Anolis tolimensis* that inhabits the lower strata of vegetation. These two species differ in their body plan, not only in body shape and external morphological features, but also in the skeleton and appendicular musculature. The results highlight the muscle and bone specializations associated with the use of habitat in this genus, such as the presence of more robust bones to enlarge the surface of muscle insertion, the thickening and loss of carpal parts, thickening of tendons associated with the manus, and greater development of muscle mass in the forelimbs by *A. heterodermus* with respect to *A. tolimensis*. These differences are related to the use of the microhabitat and the locomotor style of each species.

## INTRODUCTION

The *Anolis* lizards have been a model of study in ecomorphology in the Caribbean islands since the species that share the same type of microhabitat also share similar morphological features (Losos, 1992; Beuttell & Losos, 1999). Thus, the ecomorphs describe the correlated evolution of the morphological and ecological features in species that occupy the same space with a diversity of microhabitats (Williams, 1972; Poe & Anderson, 2019). Although equivalent processes have been observed in continental lizards, little has been studied about the morphology of continental *Anolis* and the various specieś relation to ecology, habitat use and the interactions that regulate these processes. In 2016, Moreno-Arias and Calderón-Espinosa described morphological diversity in 51 *Anolis* species from northern South America, defining as a result ten different morphological groups (morphotypes) determined by similar characteristics such as body shape, size, proportion of limbs and subdigital lamellae, possibly originated through an adaptive radiation pattern similar to that observed in other studies in the Caribbean islands (Losos & Miles, 2002; Losos, 2009). Later studies also support that adaptive radiation could explain the origin of similar *Anolis* morphotypes of continental and insular environments (Poe et al. 2018; Poe & Anderson, 2019).

Colombia has the greatest *Anolis* diversity in the world with 78 described species (Uetz & Hošek, 2019), among which we find species with different life styles. *Anolis heterodermus* Duméril 1851 is a strictly arboreal species that uses the thin branches of vegetation and lives in ecosystems of scrub and Andean forests (Miyata, 1983; Vargas-Ramírez & Moreno-Arias, 2014), and is classified as morphotype MT4 (Moreno-Arias & Calderón-Espinosa, 2016); along with other strictly arboreal continental lizards of compressed bodies and short limbs that correspond ecologically to the ecomorph “twig” in studies carried out on insular lizards (Miyata, 1983; Torres-Carvajal et al. 2010; Vargas-Ramírez & Moreno-Arias, 2014). On the other hand, *Anolis tolimensis* Werner 1916 is a species that inhabits the lower strata of Andean forest, using mainly the soil and the lower parts of trunks and branches (Ardila-Marín et al. 2008), and it is classified as morphotype MT2, along with other small lizards with a cylindrical or depressed body and long limbs, corresponding ecologically to the “grass-bush” ecomorph in studies carried out in the Caribbean islands (Moreno-Arias & Calderón-Espinosa, 2016). Consequently, these species are a model for comparative functional studies, given the morphological differences in the shape of the bodies, the sizes and proportions of their limbs as well as their habitat use. It is likely that these variations are also represented in their musculoskeletal characteristics and, therefore, in their locomotor performance.

In *Anolis*, as in other lizards, the correlation between limbs dimensions, the substrate and the locomotion characteristics of species has been analyzed; and a strong tendency of the organisms is recognized with related corporal plans using a habitat of equivalent forms (Losos & Sinervo, 1989; Losos & Irschick, 1996; Losos et al. 1997; Irschick & Losos, 1998, 1999; Vanhooydonck et al. 2005, 2006b). Herrel et al. (2008) analyzed the relationship between the characteristics of the appendicular skeleton and the locomotor style of *Anolis valencienni* and *Anolis sagrei* from the Caribbean islands. These species are assigned to different ecomorphs, and the authors concluded that there are differences in the musculature and the bone elements associated with the locomotor performance of each species. Likewise, studies in continental *Anolis* species support the relationship between morphological traits and locomotor performance (Velasco & Herrel, 2007; Moreno-Arias, 2014).

Therefore, taking into account the context of the morphological diversity in continental *Anolis,* we set the following objectives: (1) characterize the appendicular morphology of *A. heterodermus* and *A. tolimensis*; (2) evaluate the differences between the locomotor style and performance of these two species; (3) compare the morphology and locomotor performance between the continental *Anolis* species belonging to the MT4 and MT2 morphotypes with the insular species belonging to the “twig” and “grass-bush” ecomorphs.

## METHODS

We obtained specimens of *Anolis heterodermus* and *Anolis tolimensis* from the Reptile Collection of the Instituto de Ciencias Naturales of the National University of Colombia, Bogotá. For each species the musculature was described. We dissected 6 individuals of *A. heterodermus* (ICN 6248, 6250, 6251, 6256, ICN-R 13170, 13171) and 6 individuals of *A. tolimensis* (ICN 12836, ICN-R 12831, 12832, 12833, 12834, 12835). We identified each muscle of the anterior and posterior limbs and recorded its origin, insertion and the presence of tendons, following the nomenclature of Snyder (1954), Zaaf et al. (1999) and Herrel et al. (2008). Additionally, we classified the muscles into functional groups following Herrel et al. (2008). We extracted the muscles and stored them in vials with 70% alcohol, and we weighed them by functional groups using a PRECISA XB220A ± 0.01g analytical balance. The muscles for *A. heterodermus* were recorded in detail in **Tables 1** and **2**, and we listed the differences with *A. tolimensis*.

**Table 1.** Forelimb musculature. It shows the muscular description of forelimb based on *Anolis heterodermus* with muscle name, origin, insertion and assigned functional group. (**FG:** assigned functional group; **E&We:** elbow and wrist extensors; **Ee:** elbow extensors; **Ef**: elbow flexors; **Habd:** humerus abductors; **Hadd:** humerus adductors; **Hp:** humerus protractors; **Hr:** humerus retractors; **Wf:** wrist flexors; **R-Uro:** radio-ulnar rotation; **W&Df:** wrist and digits flexors; **We:** wrist extension).

| MUSCLE | ORIGIN | INSERTION | FG |
| --- | --- | --- | --- |
| *M. Trapezius* | It originates in mid-dorsal cervical connective tissue and neural spine of the thoracic vertebrae, superior to *M. Latissimus dorsi*. | The fibers run obliquely and converge in the posterior region of the proximal epiphysis of the humerus. | Hr |
| *M. Pectoralis pars superficialis* | Anterior aspect of the last sternal rib and half of the sternum longitudinally almost to the upper edge. | Through a fibrous tendon in the anterior proximal side of the humeral epiphysis in the humeral tubercle. | Hr |
| *M. Pectoralis pars profundus* | Ventromedial side of the sternum and interclavícula. | The fibers are inserted in the proximal region of the humeral tubercle. | Hadd |
| *M. Coracohumeralis anterior* | Cranioventral face of the coracoid. | Medial part of the humeral tubercle. | Hp |
| *M. Biceps II* | Cranioventral region of the humerus, anterior to the humeral tubercle. | Fibers converge with *M. Biceps I* and insert part in aponeurosis and part fleshy in the proximal face of ulna and radio. | Ef |
| *M. Biceps I* | Long and thin tendon in the medioventral face of the coracoid. | Fibers converge with *M. Biceps II* and insert part in aponeurosis and part fleshy in the proximal face of ulna and radio. | Ef |
| *M. Coracobrachialis longus* | By short tendon on the posterior face of the coracoid. | In the ventral aspect of the humerus, near the elbow. | Hadd |
| *M. Coracobrachialis brevis* | Along the posterior half of the ventral aspect of the coracoid. | Along the proximal 50% of the humerus. | Hadd |
| *M. Clavodeltoideus superficialis* | Ventral face of the interclavícula and posteroventral of the clavicle. | They run obliquely and are inserted in the cranial aspect of the humerus, proximal to the deltopectoral tubercle. | Hp |
| *M. Clavodeltoideus profundus* | Ventral part of the interclavícula. | Run a posterior between interclavícula and scapula; it is inserted proximally on the dorsocranial side of the humerus. | Habd |
| *M. Latissimus dorsi* | Cervical mid-dorsal connective tissue and neural spine of the thoracic vertebrae. | The fibers run anteroventrally and are inserted along the proximal dorsocaudal face of the humerus through a short and thin tendon. | Hr |
| *M. Scapulodeltoideus anterior* | At the junction between scapula and suprascapula, and on the medioventral face of the scapula. | Short tendon on the dorsal side of the humerus, prior to insertion of *M. Scapulodeltoideus posterior*. | Habd |
| *M. Scapulodeltoideus posterior* | External face of suprascapula. | Proximally on the dorsal side of humerus, near the humeral tubercle. | Habd |
| *M. T. b. pars humeralis anterior* | Cranial aspect of the humerus. | All converge near the elbow and insert into a thin tendon that rotates around the elbow and inserts into the proximal face of the ulna. | Ee |
| *M. T. b. pars humeralis posterior* | Cranial aspect of the humerus. |  |  |
| *M. T. b. pars scapulo-humeralis* | Scapulocoracoid ligament, some fibers from the caudal region of the junction with the humerus. |  |  |
| *M. T. b. pars scapularis* | Thin tendon on the lateral part of the scapular base. |  |  |
| *M. Epitrocleoanconus* | Short tendon on the ventral side of the distal aspect of the humerus. | Run down the ulna and insert along the first quarter of the ventral side of the ulna. | R-Uro |
| *M. Flexor carpi ulnaris* | Short tendon in ventral part of the most distal aspect of the humerus. | It is separated into two parts: the Lateral, is inserted in the ulna; while the medial, is inserted along the distal aspect of the ulna. | Lateral:<br>Wf;<br>Medial:<br>Ef |
| *M. Flexor digitorum longus pars ulnaris* | Two bodies that originate jointly on the distal face of the humerus in a short tendon. | The two bodies meet halfway and converge on the wrist; they are inserted through a long and thin tendon, which is divided and inserted into the distal phalanges of fingers II, III and IV. | W&Df |
| *M. Flexor digitorum longus pars radialis* | Distal epiphysis of the humerus. | Run between the radio and the ulna; is inserted with a long tendon in distal phalanges of fingers III and IV. |  |
| *M. Flexor digitorum longus pars profundus* | Ventral region of the ulna. | In the wrist by conspicuous tendon that divides and inserts into distal phalanges of fingers I, II and III. |  |
| *M. Flexor carpi radialis* | Dorsolateral surface of the distal epiphysis of the humerus. | Run along the radius and insert into the distal region of the same. | Ef |
| *M. Pronator teres* | Short tendon in ventral part of the distal aspect of the humerus. | Fleshy on the proximal fourth of the radius. | R-Uro |
| *M. Pronator accessorius* | Along the proximal two thirds of the ulna. | Short tendon in the middle third of the radius. | R-Uro |
| *M. Extensor carpi radialis* | Short tendon in the distal aspect of the humerus. | It runs along the radius and inserts in its entire dorsal region. | Ee |
| *M. Abductor longis pollicis* | Along the distal third of the ulna. | It becomes narrow towards the insertion, it is inserted tendinously in the dorsal-distal aspect of the first metacarpal in the finger I. | We |
| *M. Extensor carpi ulnaris* | Short tendon in distal epiphysis of the humerus. | Run along the ulna, the fibers insert part fleshy and part by tendon with *M. Flexor carpi ulnaris* in the distal part of the ulna; another tendon runs towards the lateral aspect of metacarpal V. | E&We |
| *M. Extensor digitorum longus pars superficialis* | Short tendon in the distal aspect of the humerus in conjunction with the *M. Extensor carpi radialis*. | Both muscles run parallel to each other through the first third of their length. It is inserted in the dorsal side of metacarpal V. | We |
| *M. Extensor digitorum longus pars profundus* | Short tendon in the distal aspect of the humerus. | Runs next to *M. Extensor carpi radialis* and inserts into the dorsal region of metacarpals II and III. | We |
| *M. Scapulohumeralis superficialis* | Cranial aspect of the ventral part of the suprascapula and the dorsal aspect of the scapula. | Proximally in the caudal aspect of the humerus. | Habd |
| *M. Scapulohumeralis profundus* | Caudal face of the scapula. | Proximal-dorsal region of the humerus. | Habd |
| *M. Coracobrachialis posterior* | Ventral surface of the coracoid, posterior to the coracoidal fenestra. | Proximally in the ventral aspect of the humerus, caudal to the epiphysis of the humerus. | Hadd |
| *M. Supracoracoideus* | Anterior dorsal side of the coracoid. | The proximal-dorsal aspect of the humerus. | Hr |
| *M. Extensores digitorum breves* | Set of muscles, all originate in the dorsal region of the ulna. | In metacarpals II, III and IV. The last one runs towards the base of the first phalange of the finger V. | We |
| *M. Pronator profundus* | In two-thirds of the ulna. | In the two thirds farthest from the radius; a thin and inconspicuous tendon accompanies the fibers throughout the entire muscle. | R-Uro |

**Table 2.** Hindlimb musculature. It shows the muscular description of hindlimb based on *Anolis heterodermus* with muscle name, origin, insertion and assigned functional group. (**FG:** assigned functional group; **Ae:** ankle extensors; **Ae&(t)f:** ankle extensors and toe flexors; **Af:** ankle flexors; **Fabd:** femur abductors; **Fadd:** femur adductors; **Fp:** femur protractors; **Fr:** femur retractors; **Ke:** knee extensors; **Kf:** knee flexors; **TFro:** tibio-fibular rotation).

| MUSCLE | ORIGIN | INSERTION | FG |
| --- | --- | --- | --- |
| *M. Transversus perinei* | Posterior ventral region of the ischium | Fleshy in the posterior region of the femur, from the proximal epiphysis to the middle of the diaphysis. | Fabd |
| *M. Puboischiotibialis* | Ventral region of the lateral pubispadic ligament, the ventrolateral side of the ischium and the ilioisquiadlic ligament. | The fibers converge in the ventro-medial cranial region of the tibia part fleshy and part with tendon. | Kf+Fadd |
| *M. Pubofibularis* | Aponeurosis comunis. | It crosses the *M. Adductor femoris* and inserts with a short and conspicuous tendon together with *M. Ilioischiofibularis* in the cranial region of the fibula | Fadd |
| *Tensor aponeurosis communis* | Aponeurosis comunis. | Cranioventral region of the femoral epiphysis. | Fp |
| *M. Ischiofemoralis posterior* | Ventral posterolateral side of the ischium; some fibers in the anterolateral part of the ischium. | Dorsocaudal region of the femoral epiphysis. | Fr |
| *M. Pubofemoralis pars ventralis* | Part on the ventral surface of the pubis and part from the puboisquiadlic ligament. | Ventrally in the proximal region of the trochanter. | Fadd |
| *M. Ischiofemoralis anterior* | Anterolateral region of the ischium (cartilaginous), the medial puboisquiadlic ligament and the mid-ventral edge of the pubis. | Ventral region of the base of the trochanter. | Fadd |
| *M. Flexor tibialis externus* | Ventral region of the ilioisquiadlic ligament. | With short aponeurosis in the ventral region of the tibial epiphysis. | Fadd |
| *M. Flexor tibialis internus* | Ilioisquiadlic ligament. | Short tendon in the cranial ventral region of the tibia. | Kf+Fadd |
| *M. Caudofemoralis longus* | Ventral process, ventral aspect of the vertebral body and the ventral aspect of the transverse process of the caudal vertebrae 2-9. | In the cranial region of the femur by means of a short and thin tendon; an accessory tendon divides into the middle of the main tendon and runs through the tibia where it is inserted distal to the knee joint. | Fr |
| *M. Caudofemoralis brevis* | Ventral region of the vertebral body and in the transverse process of the caudal vertebrae 1-4. | Ilioisquiadlic ligament. | Other hindlimb |
| *M. Adductor femoris* | The proximal fibers in the caudal region of the lateral puboisquiatric ligament; the intermediate fibers in the ventral aspect of the ischium; caudal fibers lateral to the caudal side of the ischium; a small group of fibers in the ilioisquiatric ligament. | Along the distal three quarters of the femur. | Fadd |
| *M. Ilioischiofibularis* | Ilioisquiatric ligament and ilium. | In the cranial aspect of the fibula with conspicuous tendon. | Fadd |
| *M. A. pars dorsalis* | In large and conspicuous aponeurosis along the first ascending half of the ilium. | In the proximal aspect of the tibia through a short aponeurosis. | Ke |
| *M. A. pars ventralis* | With short aponeurosis at the base of the pubis and more proximal aspect of the trochanter. | In the proximal aspect of the tibia through a short aponeurosis. | Ke |
| *M. Femorotibialis ventralis* | Along the distal two thirds of the femur. | With a thin tendon in the cranial region of the tibia. | Ke |
| *M. Femorotibialis dorsalis* | Dorsal region of the femur. | With short tendon in the dorsolateral region of the tibia. | Ke |
| *M. Pubofemoralis pars dorsalis ext.* | Dorsocranial region of the pubis. | Aponeurosis comunis. | Fp |
| *M. Pubofemoralis pars dorsalis int.* | Dorsocranial region of the pubis. | Femur, distal to the trochanter. | Fp |
| *M. Ischiofemoralis dorsalis anterior* | Dorsal region of the ischium. | In the cranial aspect of the femur, distal to the trochanter. | Fp |
| *M. Iliofibularis* | Base of the ilium, anterior to the posterior ascending process. | Fibula, deep to the dorsal part of the *M. Gastrocnemius pars fibularis major*. | Kf |
| *M. Iliofemoralis* | In the anterior ventrolateral region of the ilium | Proximally in caudal aspect of a femur at the level of insertion of *M. Caudofemoralis longus*. | Fabd |
| *M. Ilioischiotibialis* | Dorsolateral aspect of the ilioisquiatric ligament. | The first part is inserted proximally on the ventromedial side of the tibia; the other part proximally on the ventrolateral side of the tibia. It has a clear tendon in the insertion, it is divided at the level of *M. G. pars fibularis major*; the first part of the tendon is inserted proximally on the ventromedial side of the tibia; the other part runs through the tibial part of the *M. G. pars fibularis major* and inserts proximally on the ventrolateral side of the tibia. | Kf |
| *M. Tibialis anterior* | Two bodies, the first in the anterior aspect of the tibia; the second in the ventral aspect of the tibia (fleshy). This muscle is much thicker at the origin in *A. heterodermus* with respect to *A. tolimensis*. | Lateral aspect of the first metatarsal. | Af |
| *M. Extensor digitorum longus* | With long and thin tendon in the fibular region of the femur. | Dorsal side of the third metatarsal. | Af |
| *M. Gastrocnemius pars fibularis major* | Thin tendon on fibular side of thoracic dorsal tubercle. | On the first phalanx of fingers IV and V. | Ae |
| *M. Gastrocnemius pars fibularis minor* | Thin tendon on fibular side of thoracic dorsal tubercle. | On the first phalanx of finger IV. | Ae |
| *M. Gastrocnemius pars profundus* | Small and laminar, tibial side of the distal part of the femur. | Short tendon that crosses to the other side, runs under the plantar aponeurosis and inserts medially at the level of the fifth metatarsal. | Ae |
| *M. Flexor digitorum communis* | Two parts: the tibial part fleshy in the proximal third along the interior aspect of the tibia and fibula; the fibular part in the proximal third along the inside of the fibula. | Tibial part in distal phalanx of fingers I-IV; fibular part in the distal phalanx of the V toe. The two bodies converge on a tendon that wraps around the ankle. | Ae&(t)f |
| *M. Extensor ossi metatarsi hallucis* | Ventral region of the distal two thirds of the fibula | Short tendon in the dorsal aspect of the tibial side of the astragalocalcaneum. | Ae |
| *M. Peroneus brevis* | Two thirds distal from the cranial edge of the fibula. | Posterodorsal side of the fifth metatarsal. | Ae |
| *M. Popliteus* | Medial side of the most proximal region of the fibula | On the medial side of the fifth proximal of the tibia. | TFro |
| *M. Peroneus longus* | Long and thin tendon on the fibular side of the femur. | It is wrapped around the knee and inserted into the ventral side of the fifth metatarsal. | Ae |
| *M. Iliofemoralis posterior* | Posterior region of the ilium (accessory to the ilioisquiatric ligament) and from the ventral aspect of the first caudal vertebra. | Conspicuous tendon on the femur. | Fabd |
| *M. Ischiofemoralis dorsalis posterior* | Dorsocaudal side of the ischium. | Dorsocaudal side of the femur. | Fabd |
| *M. Pronator profundus* | Fleshy in the distal quarter on the medial aspect of the fibula. | It runs obliquely and ventrally and inserts on the mesial side of the fifth distal of the tibia. | TFro |

The description of the appendicular skeleton of *A. heterodermus* and *A. tolimensis* was based on two adult specimens of each species (ICN-R-6287, 6251, 13036, 13027). We prepared the skeletons using the technique of differential staining for bone and cartilage from Wassersug (1976). The osteological description follows the Krause nomenclature (1989), and the description of the sesamoids follows Jerez et al. (2010). Observations were made with a NIKON C-LEDS stereoscope, and specimens were photographed with a LEICA M125 motorized stereomicroscope.

For the locomotor performance tests we obtained live specimens of the two species. We captured 30 adult specimens of *Anolis heterodermus* (17 males: SVL = 65.7 mm average ± 4.80 mm SD and 13 females: SVL = 64.5 ± 5.69 mm) from the municipality of Tabio, Cundinamarca (Colombia) between April and May 2016. The individuals were transported to the Evolutionary Ecology Lab of the National University of Colombia in Bogotá, where they were kept in terrariums with branches and plant material at room temperature (17-19 ° C), thanks to the temperature in Bogotá is equivalent to the operational temperature of the species in its natural environment (Méndez-Galeano & Calderón-Espinosa, 2017). Individuals were fed twice a day with small insects and the terrariums were sprayed with water twice a day to keep the environment humid. Once the performance tests were performed, the individuals were released at the same capture site.

Likewise, we captured 30 adult individuals of *A. tolimensis* (11 males: SVL = 47.5 ± 2.28 mm and 19 females: SVL = 50.2 ± 2.22 mm) in the municipality of Silvania, Cundinamarca (Colombia) between May and July of 2017. These were temporarily stored in wet canvas bags with vegetation, while the performance tests were carried out at the capture site; and they were immediately released at the same site.

For all individuals of the two species that were tested for performance we recorded sex (presence of hemipenia, body size), weight (balance OHAUS CL201 ± 0.1g), body dimensions (snout-vent length [SVL] and tail length [Lco], forelimb length: humerus [Lbr], radius-ulna [Labr], metacarpus and length of the longest digit not including the claw; hindlimb length: femur [Lm], tibia [Lp], metacarpus [Ldp] and length of the longest toe without including the claw). We used a digital electronic caliper (REDLINE M ± 0.02 mm) and always took the lengths of the limbs of the right side.

For the performance tests we filmed each individual from the side view using a NIKON D3300 camera at 60 frames per second (60p) as the animals ran on two cylindric platforms of different diameters (10 mm and 80 mm) that were 2m long and arranged at an inclination of 45°. The animals were encouraged to run as fast as possible clapping behind them or touching their tails with a thin paintbrush. At least three sequences per individual were obtained and analyzed for each platform.

For all the analyses we performed ANOVA and Kruskal-Wallis statistical tests according to the parametric or non-parametric nature of the data. Following the methodology proposed by Sokal & Rohlf (1995) we used the Shapiro-Wilks test to evaluate normality and the Levene test for homoscedasticity.

Initially we evaluated differences in snout-vent length (SVL) and other body measurements (LCo, Lbr, Labr, Lm, Lp and Ldp) between species. We also analyzed the differences between the proportions of the hindlimbs with respect to the forelimbs (Lep/Lea) in total length and in each of its components (arm-thigh and forearm-leg). This analysis was done with a correction for size (SVL) in order to eliminate the error produced by the difference in the size of the species.

In order to establish which were the most important muscle groups during the locomotion of each species and their differences, we did an analysis of variance from the weights recorded by the functional groups of each species, corrected with the total weight of each.

We analyzed the videos using the software TRACKER (version 4.10.0; Brown, 2017) examining from the moment the lizard starts to move until it leaves the frame of the scene (pixel error margin <20%). Based on this data, we quantified the average speed of movement obtained by analyzing the position of the tip of the snout (distance traveled in the frame) with respect to time. We also quantified the length of the step, that is, the distance that the body moves forward during the support of the hindlimb and finally the step frequency, which is the number of steps per second. These variables were obtained by analyzing the frames in which the right foot was in contact with the substrate.

To analyze the locomotor performance, a correlation analysis was performed (Pearson test for the parametric samples and Spearman test for the non-parametric samples) taking in account the locomotion variables with respect to the size of the individuals, in order to evaluate whether these variables are determined by size. From this result, we corrected the data with the SVL values of each individual. We carried out a bidirectional multivariate analysis of nonparametric variance (two-way PERMANOVA (Anderson, 2001)) to establish the level of interaction of the species and platforms on the locomotion variables evaluated. In addition, we did a univariate comparison using the ANOVA and Kruskal-Wallis tests to establish the influence of the species and the surface among the locomotion variables studied. Following the methodology proposed by Herrel et al. (2008), only the movements made after the third step in each event were considered, that corresponds to the acceleration stage. We performed all the statistical tests using the software Past 3.15 (Hammer et al. 2001).

## RESULTS

### Morphometrics

*Anolis heterodermus* is a medium-sized lizard (SVL = 65.78 ± 4.96 mm) with a head length of almost 23% of its body (LaCa = 22.14 ± 0.72 mm). Its tail is long (LCo = 85.59 ± 9.63 mm) in relation to the length of its body. It is not sexually dimorphic in terms of size (Moreno-Arias, 2014). The species has a compressed and robust body. The forelimbs are longer and more robust (26.96 ± 3.82 mm) than the hindlimbs (22.71 ± 2.94 mm) as can be seen in **Figure 1-A**.

**Figure 1.**
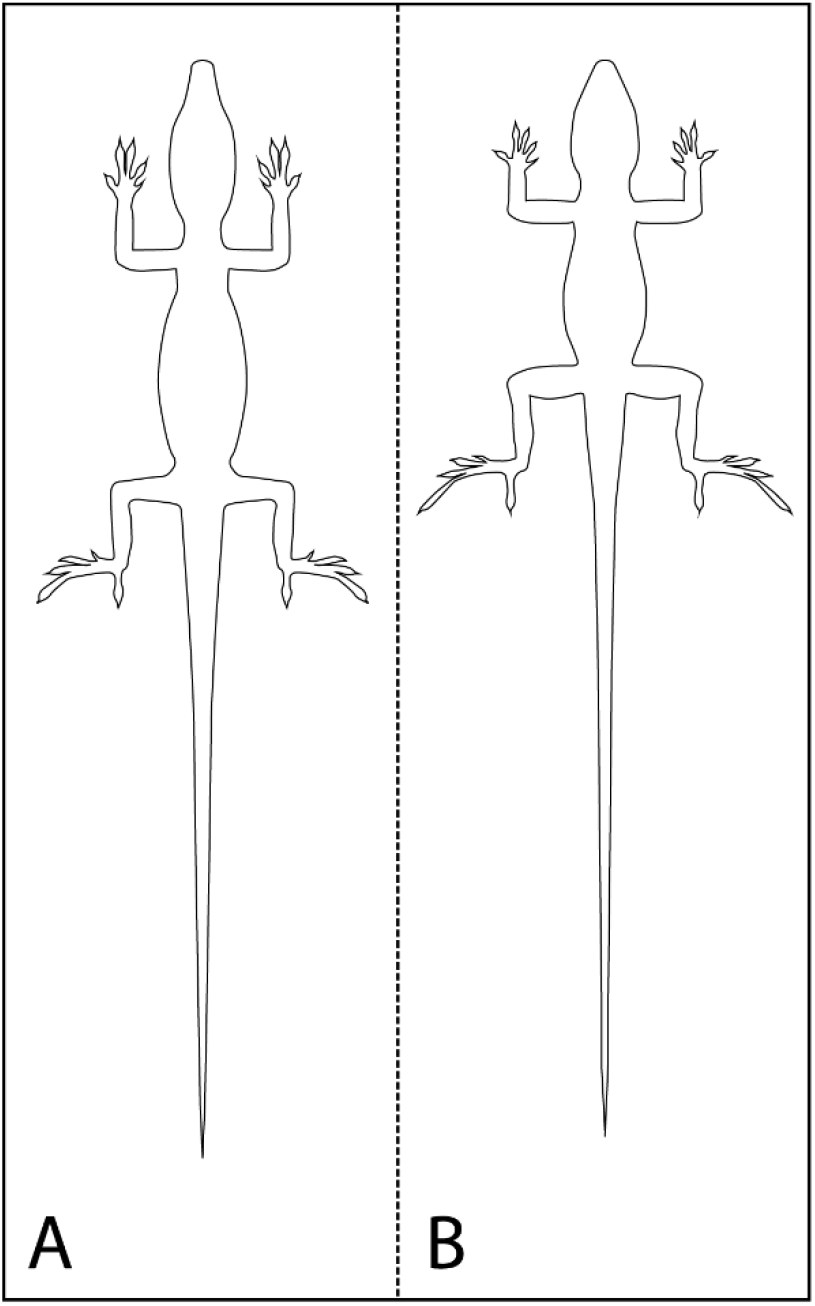
Body shape of *Anolis heterodermus* and *Anolis tolimensis*. It shows the body shape of *A. heterodermus* (A) and *A. tolimensis* (B).

*Anolis tolimensis* is a small lizard (SVL = 49.36 ± 2.56 mm) with a head length of almost 26% of its body (LaCa = 13.33 ± 0.84 mm). It has a very long tail (LCo = 97.11 ± 17.42 mm), twice the length of the body. It is sexually dimorphic with females larger than males (Ardila-Marín et al. 2008). The males have a thin and compressed body, while the females are robust and have a cylindrical body. This species has short and thin forelimbs (22.06 ± 1.38 mm) as compared to the hindlimbs that are approximately twice the length (40.70 ± 2.10 mm) of the forelimbs as shown in **Figure 1-B**.

*Anolis heterodermus* is a significantly larger species than *A. tolimensis* (Kruskal-Wallis: H1.56 = 39.96, P = 2.581E-10). In general, the two species varied significantly in tail length (Kruskal-Wallis: H1.54 = 40.79, P = 1.695E-10), forearm length (Kruskal-Wallis: H1.56 = 5.999, P = 0.01431), thigh length (ANOVA: F1.56 = 105.9, P = 1.61E-14), leg length (Kruskal-Wallis: H1.56 = 41.15, P = 1.409E-10) and foot length (Kruskal-Wallis: H1.56 = 42.36, P = 7.605E-11). The exception was the arm length (Kruskal-Wallis: H1.56 = 1.185, P = 0.2763) which is similar in the two species.

An analysis of the differences between the proportions of the limbs with respect to the body of each species shows that *A. tolimensis* has a Lep / Lea ratio significantly higher than that of *A. heterodermus* (Kruskal-Wallis: H1.56 = 36.31, P = 1.68E-09), both in the total length of the limb and in each of its components (Thigh-Arm: ANOVA: F1.56 = 23.07, P = 1.21E-05 and Leg-Forearm: Kruskal-Wallis: H1.56 = 39.96, P = 2.59E-10). Therefore, both limbs of *A. tolimensis* are longer proportionally than *A. heterodermus*.

### Appendicular skeleton

#### Anolis heterodermus

*Pectoral girdle:* it has a complete girdle (**Figure 2-A, C**). The clavicle is depressed and widened in almost all its extension, only compressed in the lateral end; it extends from the midline to the acromial process region. The interclavicle, T-shaped and depressed, has the medial process very wide anteriorly, but narrow and acute posteriorly; this extends to 1/3 of the pre-sternum; the lateral processes are very wide, with the distal end straight and extend up to ¾ of the length of the clavicle, and do not contact the coracoid. The epicoracoid, laminar and narrow, closes the coracoid fenestra anteriorly, and lies on the meso-sternum posteriorly. The coracoid presents the mesocoracoid thin and long; whereas, the metacoracoid is broad and slightly concave at the base and extends anteriorly where it becomes narrower and slightly convex; also, it presents the coracoid fenestra and the coracoid foramen. The scapula is quadrangular and presents a large process on the anterior edge very broad at the base and pointed, which forms the lateral edge of the scapula-coracoid fenestra. The suprascapula is broad at the base, and dorsally widens, forming almost a semicircle. The pre-sternum is rhomboidal, broad towards the anterior region, and narrow in the posterior region, where it articulates with two pairs of ribs; the meso-sternum is formed by two thin and long bars and articulates with two pairs of ribs.

**Figure 2.**
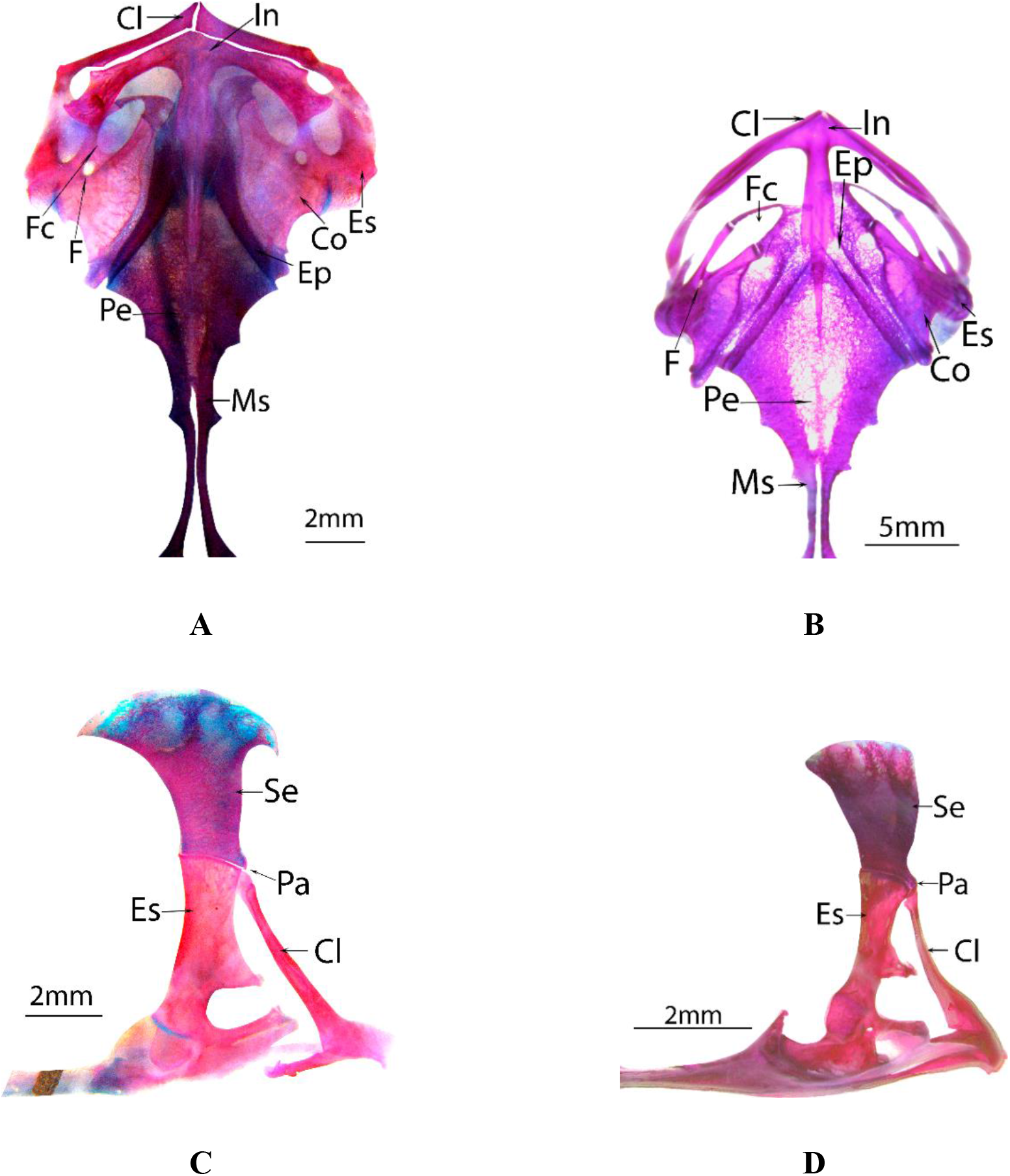
Pectoral girdle of *Anolis heterodermus* and *Anolis tolimensis*. *A. heterodermus* ventral (**A**) and lateral (**C**) view. *A. tolimensis* ventral (**B**) and lateral (**D**) view. (**Cl**: clavicula; **Co**: coracoid; **In**: interclavicula; **Ep**: epicoracoid; **Es**: scapula; **F**: coracoid foramen; **Fc**: coracoid fenestra; **Ms**: meso-sternum; **Pa**: acromial process; **Pe**: pre-sternum; **Se**: supra-scapula; bar = 2mm (A, C, D), 5mm (B)).

*Pelvic girdle:* it has a complete, wide and robust girdle (**Figure 3-A, C**). The pubis is wide and exhibits a very wide and pointed prepubic process, which extends anteriorly from the base of the pubis. The epipubis is rhomboidal and is fused to the pubis. The ischium is wide and quadrangular, exhibits the ischiadic process posteriorly, which is very wide at the base and pointed. The ischiadic symphysis is ossified. The hipoischium is thin, short and cartilaginous. The obturator fenestra is broad and cordiform. Finally, the ilium is very compressed and wide in all its extension; presents a short process at the base of the anterior edge, which is broad at the base and pointed; it is positioned at 45° with respect to the longitudinal axis and articulates with the sacral vertebrae.

**Figure 3.**
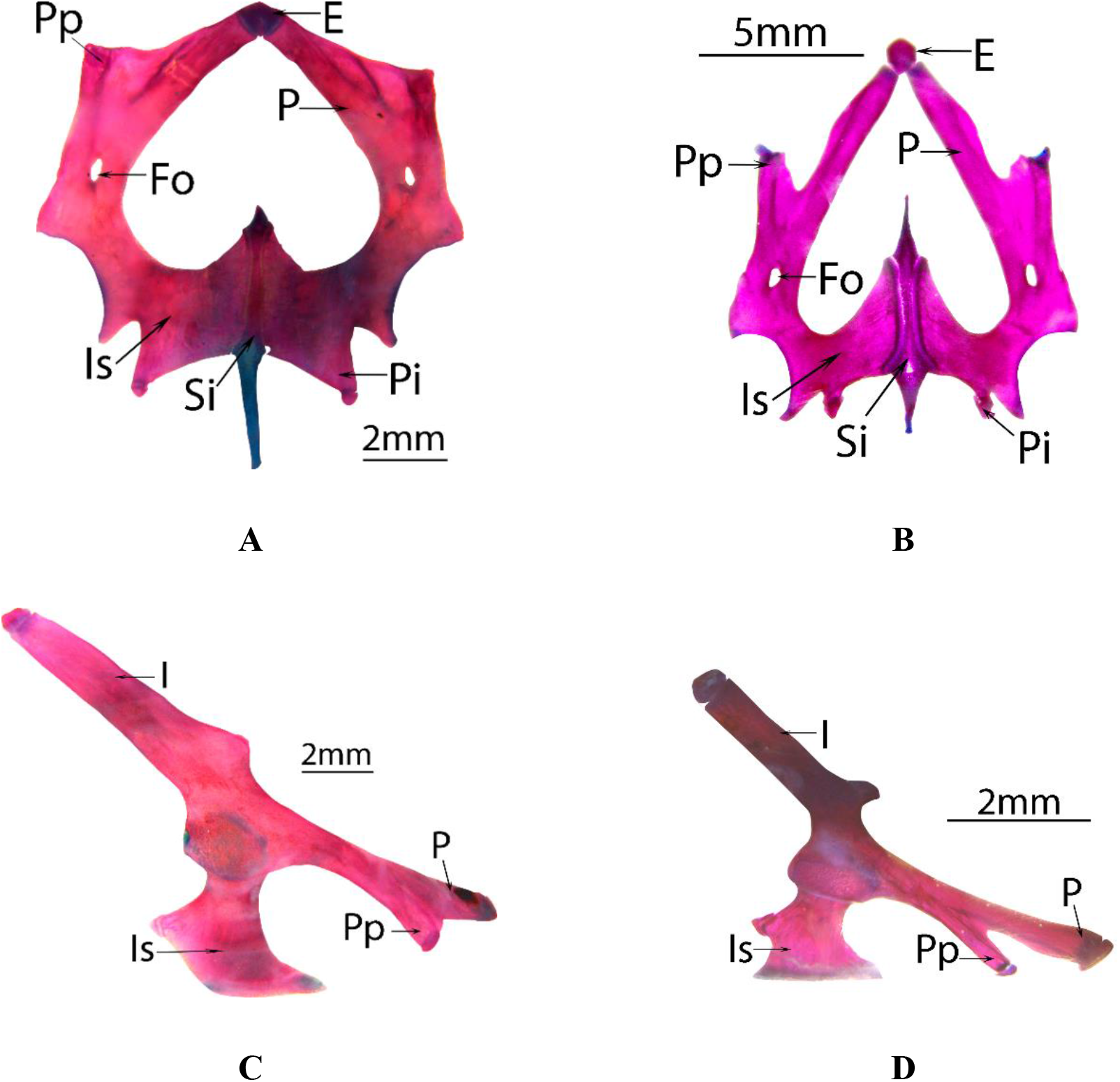
Pelvic girdle of *Anolis heterodermus* and *Anolis tolimensis.* *A. heterodermus* ventral (**A**) and lateral (**C**) view. *A. tolimensis* ventral (**B**) and lateral (**D**) view. (**E:** epipubis; **Fo:** obturator fenestrae; **I:** ilium; **Is:** ischium; **P:** pubis; **Pi:** ischiatic process; **Pp:** prepubic process; **Si:** ischiatic symphysis; bar = 2mm (A, C, D), 5mm (B)).

*Forelimb:* constituted by humerus, radio-ulna, carpals, metacarpals and five digits (**Figure 4-A**). The carpal elements of the proximal series are very robust, with a quadrangular ulnar bone, an almost rectangular radial and a triangular central. Regarding to the distal series, only carpals 2 to 5 are observed, carpal 1 being absent; the carpal 4 stands out, since it is longer and wider than the other carpals. The diaphyses of the metatarsals are thin, while the proximal epiphyses are widened, especially the one of metacarpal I; metacarpal III is the longest, and decreases in size III, II, IV, I, V. The phalangeal formula is 2-3-4-5-3. The terminal phalange is a claw, with a ventral and proximal process, which is wide, low and sharp. The sesamoid elements are the ulnar patella, the distal supraphalangial sesamoids, the palmar, the pisiform, and the sesamoid anterior to the pisiform. In general, the bones of the stylopodium and the zeugopodium are robust and thick.

**Figure 4.**
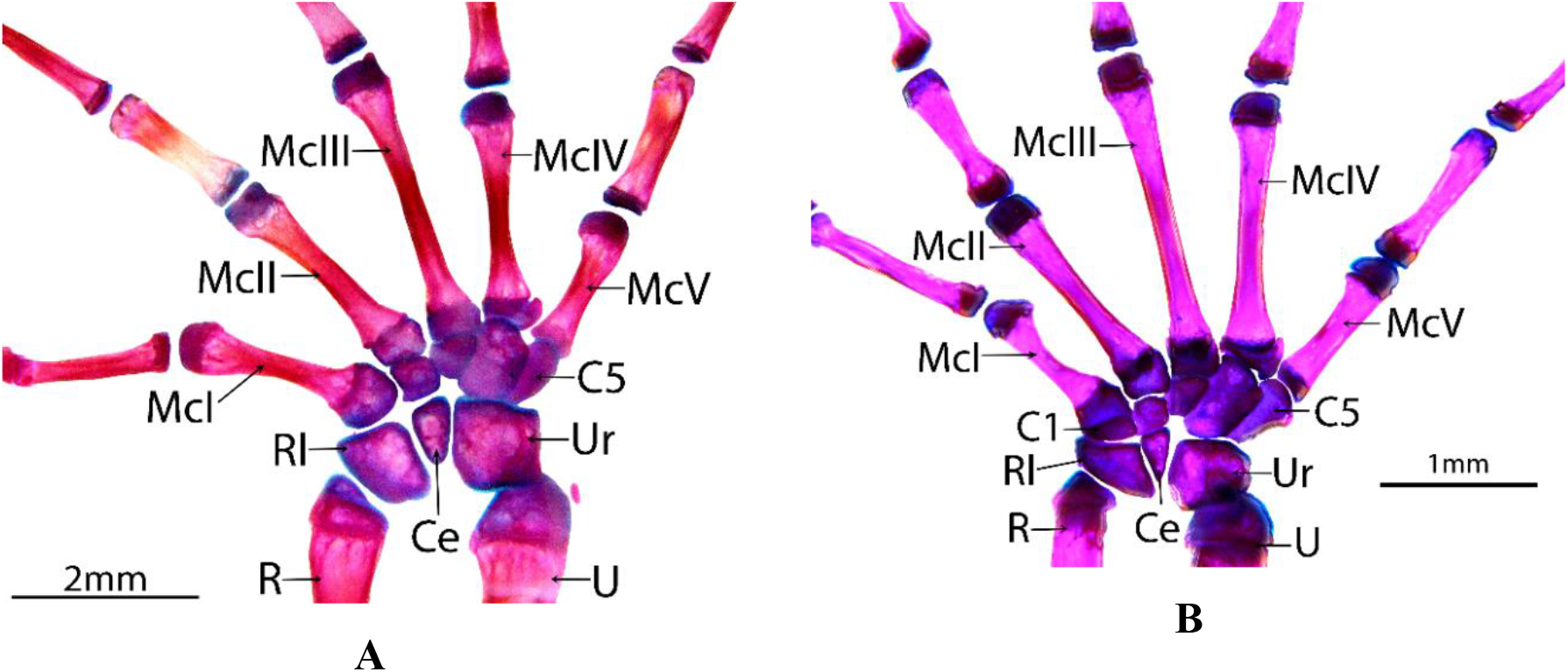
Carpus and metacarpus of *Anolis heterodermus* and *Anolis tolimensis.* Dorsal view *A. heterodermus* (**A**) and *A. tolimensis* (**B**). (**C1:** carpal 1; **C5:** carpal 5; **Ce:** central; **McI-McV:** metacarpals I to V; **R:** radius; **Rl:** radial; **U:** ulna; **Ur:** ulnar; bar = 2mm (A), 1mm (B)).

*Hindlimb:* is conformed by femur, tibia-fibula, tarsus, metatarsus and five toes **(Figure 5-A, C**). The tarsus has the proximal tarsal with a conspicuous dorsal concavity, and the distal tarsals III and IV. In the metatarsals, the diaphyses are thin and the epiphyses are slightly widened; metatarsal IV is the longest of the series, which decreases in sequence IV, III, II, I, V. The phalangeal formula is 2-3-4-5-4. The terminal phalange is a claw, with a short proximal process and pointed on the lower edge. The sesamoid elements are the tibial lunula, the dorsal tarsal sesamoid and the distal supraphalangeals. In general, the bones of the zeugopodium and stylopodium are thick and robust, which is very noticeable in the femur that shows the proximal end of the diaphysis quite widened.

**Figure 5.**
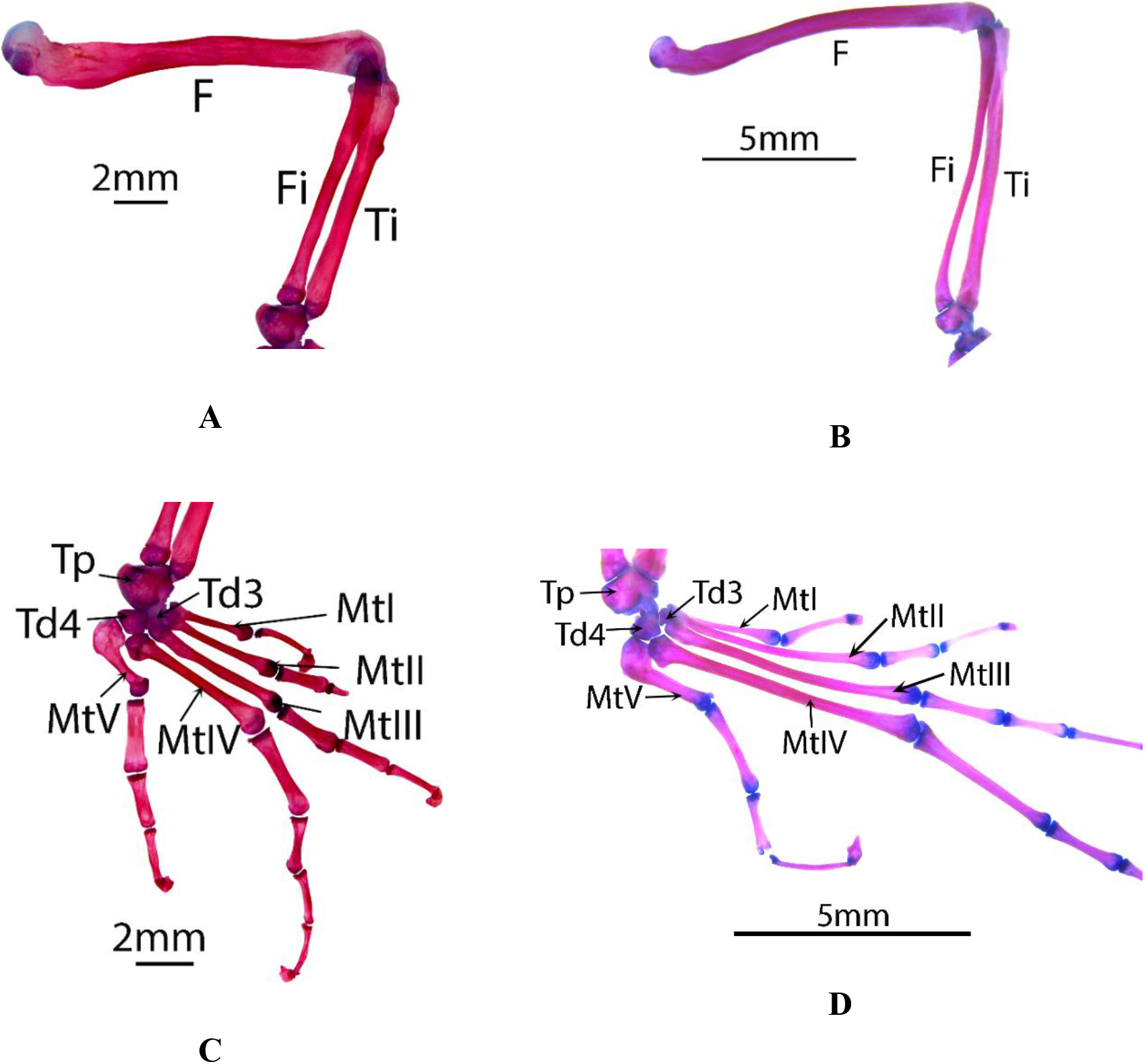
Hindlimb of *Anolis heterodermus* and *Anolis tolimensis.* *A. heterodermus* dorsal view of long bones (**A**) and dorsal view of tarsal bones (**C**). *A. tolimensis* dorsal view of long bones (**B**) and dorsal view of tarsal bones (**D**). (**F:** femur; **Fi:** fibula; **MtI- MtV:** metatarsals I to V; **Tp:** proximal tarsal; **Td3:** distal tarsal 3; **Td4:** distal tarsal 4; **Ti:** tibia; bar = 2mm (A, C), 5mm (B, D)).

#### Anolis tolimensis

*Pectoral girdle*: it has a complete girdle (**Figure 2-B, D**). The clavicle extends from the midline to the region of the acromial process; it is depressed, wide in the medial region and narrow towards the lateral ends. The T-shaped interclavicula has a wide, pointed medial process and extends to the middle region of the sternum; the lateral processes are wide, thinning towards the ends and ending in acute form, and do not contact the coracoid. The epicoracoid is laminar and narrow, lies on the sternum, surrounds the coracoid and closes anteriorly with the coracoid fenestra. The coracoid has two processes that limit the coracoid fenestra: the anterior and thin mesocoracoid, and the metacoracoid wide at the base and slightly convex anteriorly; presents the coracoid foramen. The scapula is rectangular and exhibits a process on the anterior edge, wide at the base and pointed, which forms the lateral border of the scapula-coracoid fenestra. The suprascapula is wide at the base, but dorsally it only widens slightly. The pre-sternum is rhomboidal and articulates with two pairs of ribs; The meso-sternum is constituted by two long and thin bars, which articulates with three pairs of ribs.

*Pelvic girdle*: it has a complete and narrow appearance (**Figure 3-B, D**). The pubis is narrow in the anterior middle region and wide in the posterior middle region; in this last region it presents the prepubic process wide at the base, pointed, and extending anteriorly. At the base of the pubis it encounters the obturator foramen. Anteriorly, the pubis articulates with the epipubis, that haves a hexahedral shape. The ischium is rectangular, and very wide towards the middle region; presents a short, narrow and acute sciatic process. The obturator fenestra is narrow and cordiform. The ilium is wide, compressed and presents a narrow, short and blunt process at the base and towards the front edge.

*Forelimb*: constituted by humerus, radio-ulna, carpus, metacarpus and phalanges (**Figure 4-B**). The carpus presents the radial, the ulnar, the central and the carpals 1-5; with carpals 4 and 5 large and the other small. The metapodium is made up of five metacarpals; where III is the longest of the series that decreases in sequence III, IV, II, V, I. The phalangeal formula is 2-3-4-5-3. The terminal phalange is a sharp claw, with a short and blunt ventral process. The sesamoid elements are the ulnar patella, pisiform, palmar and distal supraphalangial. In general, the bones of the stylopodium and zeugopodium are thin and gracile.

*Hindlimb:* it is complete (**Figure 5-B, D**). The tarsus presents the proximal tarsal, the distal tarsals III and IV. The metatarsal is composed of five metatarsals, where the IV is the longest of the series that decreases in sequence IV, III, II, I, V. The phalangeal formula is 2-3-4-5-4. The terminal phalange is an acute claw, with a short proximal and blunt process in the lower margin. The elements of the sesamoids are the tibial patella, the tibial lunula, the dorsal tarsal sesamoid and the supraphalangial distal. In general, the bones of the stylopodium and zeugopodium are thin, the femur has a curved appearance and the fibula is very thin with respect to the tibia.

### Muscles and distribution of muscle mass

We identified all the muscles of the anterior and posterior limbs in the two species, detailing their origin and insertion (**Tables 1, 2**). We grouped the muscles into functional groups, following the proposal of Herrel et al. (2008). Once corrected by body mass (**Table 3**), the functional groups showed that the two species do not differ significantly in the total muscle mass (Kruskal-Wallis: H1.10 = 0.9231, P = 0.3367). We observed significant differences regarding the total weight of the muscles of each limb (**Table 4**), where the hindlimb had greater weight than the forelimb in the two species (forelimb ANOVA: F1.10 = 522, P = 5.819E-10, hindlimb Kruskal-Wallis: H1.10 = 8.308, P = 0.003948). However, in *A. heterodermus* the mass of the forelimb corresponds to 83% of the mass of the hindlimb, while in *A. tolimensis* the forelimb mass is only 43% of the hindlimb.

**Table 3.** Average values and standard deviation in the functional groups muscle mass.

| **Functional group** | *Anolis heterodermus* |  | *Anolis tolimensis* |  |
| --- | --- | --- | --- | --- |
|  | **Average (mg)** | **St.dev** | **Average (mg)** | **St.dev** |
| Snout–vent length (mm) | 65,46 | 5,18 | 49,4 | 2,39 |
| Total hindlimb muscle mass | 23,93 | 3,73 | 30,70 | 4,86 |
| Total forelimb muscle mass | 19,95 | 3,63 | 13,06 | 2,77 |
| Femur protractors | 2,27 | 0,67 | 0,93 | 0,29 |
| Femur retractors | 0,41 | 0,15 | 8,12 | 1,20 |
| Femur abductors | 0,49 | 0,24 | 1,56 | 0,68 |
| Femur adductors | 7,70 | 0,53 | 6,09 | 0,68 |
| Knee flexors | 6,52 | 0,98 | 4,96 | 0,69 |
| Knee extensors | 3,22 | 0,42 | 4,34 | 0,65 |
| Ankle flexors | 0,58 | 0,11 | 0,79 | 0,11 |
| Ankle extensors | 1,85 | 0,41 | 3,07 | 0,36 |
| Other hindlimb | 0,90 | 0,21 | 0,82 | 0,20 |
| Humerus retractors | 7,06 | 0,73 | 5,54 | 0,38 |
| Humerus protractors | 0,58 | 0,22 | 0,35 | 0,14 |
| Humerus abductors | 1,49 | 0,21 | 0,77 | 0,21 |
| Humerus adductors | 1,96 | 0,27 | 1,19 | 0,16 |
| Elbow flexors | 2,58 | 0,32 | 1,43 | 0,39 |
| Elbow extensors | 2,72 | 0,41 | 1,48 | 0,27 |
| Wrist flexors | 1,08 | 0,32 | 0,81 | 0,44 |
| Wrist extensors | 1,11 | 0,37 | 0,55 | 0,24 |
| Other forelimb | 1,37 | 0,79 | 0,96 | 0,53 |
| Total muscle mass | 43,88 | 7,36 | 43,77 | 7,62 |

**Table 4.** Results of analysis of variance for the muscle mass of functional groups in *Anolis heterodermus* and *Anolis tolimensis*.

| Functional group | Normality | Homoscedasticity | F<sub>1,10</sub> | P (Same) | H<sub>1,10</sub> | P (Same) |
| --- | --- | --- | --- | --- | --- | --- |
| Total muscle mass |  | X |  |  | 0,922 | 0,3367 |
| Total hindlimb muscle mass | X |  | - | - | 8,308 | 0,003948 |
| Total forelimb muscle mass | X | X | 522 | < 0,0001 | - | - |
| Femur protractors | X | X | 0,1808 | 0,6797 | - | - |
| Femur retractors |  |  | - | - | 8,308 | 0,003948 |
| Femur abductors | X | X | 13,7 | 0,004103 | - | - |
| Femur adductors | X | X | 21,33 | 0,0009523 | - | - |
| Knee flexors |  | X | - | - | 8,308 | 0,003948 |
| Knee extensors | X | X | 12,22 | 0,00577 | - | - |
| Ankle flexors | X | X | 266,9 | < 0,0001 | - | - |
| Ankle extensors | X | X | 73,79 | < 0,0001 | - | - |
| Other hindlimb | X | X | 11,52 | 0,006841 | - | - |
| Humerus retractors | X | X | 28,69 | 0,0003207 | - | - |
| Humerus protractors | X | X | 0,9937 | 0,3423 | - | - |
| Humerus abductors | X | X | 0,5148 | 0,4895 | - | - |
| Humerus adductors | X | X | 3,97 | 0,07432 | - | - |
| Elbow flexors | X | X | 31,98 | 0,0002109 | - | - |
| Elbow extensors | X | X | 39,67 | < 0,0001 | - | - |
| Wrist flexors | X | X | 1,429 | 0,2596 | - | - |
| Wrist extensors | X | X | 0,2105 | 0,6562 | - | - |
| Other forelimb |  | X | - | - | 0,641 | 0,4159 |

Based on the average muscle mass, some functional groups stand out. The humeral retractor muscles are the heaviest in the forelimb of both species, followed by the elbow extensors and the elbow flexors. Likewise, these same functional groups are heavier in *A. heterodermus* than in *A. tolimensis*. In the hindlimb the two species differ in the functional group with a greater muscle mass. For *A. heterodermus* the femur adductors followed by the knee flexors and the knee extensors are the heaviest muscles groups of the hindlimb. But for *A. tolimensis* the heaviest muscle groups are the femur retractors, followed by the femur adductors and the knee flexors (**Table 3**).

In the forelimb there are significant differences between the two species only in some functional groups such as the humeral retractors (ANOVA: F1.10 = 28.69, P = 0.0003207), elbow flexors (ANOVA: F1.10 = 31.98, P = 0.0002109) and elbow extensors (ANOVA: F1.10 = 39.67; P = 0.00008928), which present higher mass in *A. heterodermus* than in *A tolimensis*.

The results of the analysis of variance show significant differences in all the muscle groups of the hindlimb between both species, except for the femoral protractors (ANOVA: F1.10 = 0.1808, P = 0.6797). *A. heterodermus* has greater muscle mass in the femur adductors (ANOVA: F1.10 = 21.33, P = 0.0009523), knee flexors (Kruskal-Wallis: H1.10 = 8.308, P = 0.003948) and other hindlimb muscles (ANOVA: F1.10 = 11.52, P = 0.006841), while *A. tolimensis* shows greater development in femur retractors (Kruskal-Wallis: H1.10 = 8.308, P = 0.003948), femoral abductors (ANOVA: F1.10 = 13.7, P = 0.004103), knee extensors (ANOVA: F1.10 = 12.22, P = 0.00577), ankle flexors (ANOVA: F1.10 = 266.9, P = 0.00000001536) and ankle extensors (ANOVA: F1.10 = 73.79, P = 0.000006281).

### Locomotor performance

In the applied correlation analysis, all the variables were correlated with body size (all P< 0.05). The respective corrections were made in such a way that the variance due to this factor was eliminated *a priori*. The two-way PERMANOVA showed that both the species and the platform had a significant effect on locomotor performance (Species: F1.116 = 225.01, P = 0.0001. Platform: F1.116 = 16.201; P = 0.0001); and the interaction effects were significant (Interaction: F1.116 = 10.547, P = 0.0004).

The ANOVA and Kruskal-Wallis tests showed significant differences in most of the comparisons in the variables analyzed (**Table 5**). The exception was in the comparisons of step frequency between both platforms for *A. tolimensis* (ANOVA: F1.58 = 1.878; P = 0.1758) and the step frequency between species for the wide platform (ANOVA: F1.58 = 1.099; P = 0.2988). This established that *A. tolimensis* did not vary the step frequency between different substrates, whereas *A. heterodermus* decreased this frequency on narrow substrates (**Figure 6**).

**Table 5.** Values of the univariate analysis for gait characteristics in *Anolis tolimensis* and *Anolis heterodermus* in different substrates.

|  | Statistic | Value | *p* |
| --- | --- | --- | --- |
| Between substrates for *Anolis tolimensis* |  |  |  |
| Step length | H<sub>1,58</sub> | 24,38 | < 0,0001 |
| Step frequency | F<sub>1,58</sub> | 1,878 | **0,1758** |
| Average speed | F<sub>1,58</sub> | 13,24 | 0,0005844 |
| Between substrates for *Anolis heterodermus* |  |  |  |
| Step length | F<sub>1,58</sub> | 6,721 | 0,01204 |
| Step frequency | H<sub>1,58</sub> | 11,16 | 0,0008339 |
| Average speed | H<sub>1,58</sub> | 23,23 | < 0,0001 |
| Between species in the narrow substrate |  |  |  |
| Step length | H<sub>1,58</sub> | 8,831 | 0,00296 |
| Step frequency | H<sub>1,58</sub> | 24,53 | < 0,0001 |
| Average speed | H<sub>1,58</sub> | 44,07 | < 0,0001 |
| Between species in the wide substrate |  |  |  |
| Step length | H<sub>1,58</sub> | 52,04 | < 0,0001 |
| Step frequency | F<sub>1,58</sub> | 1,099 | **0,2988** |
| Average speed | H<sub>1,58</sub> | 41,55 | < 0,0001 |

**Figure 6.**
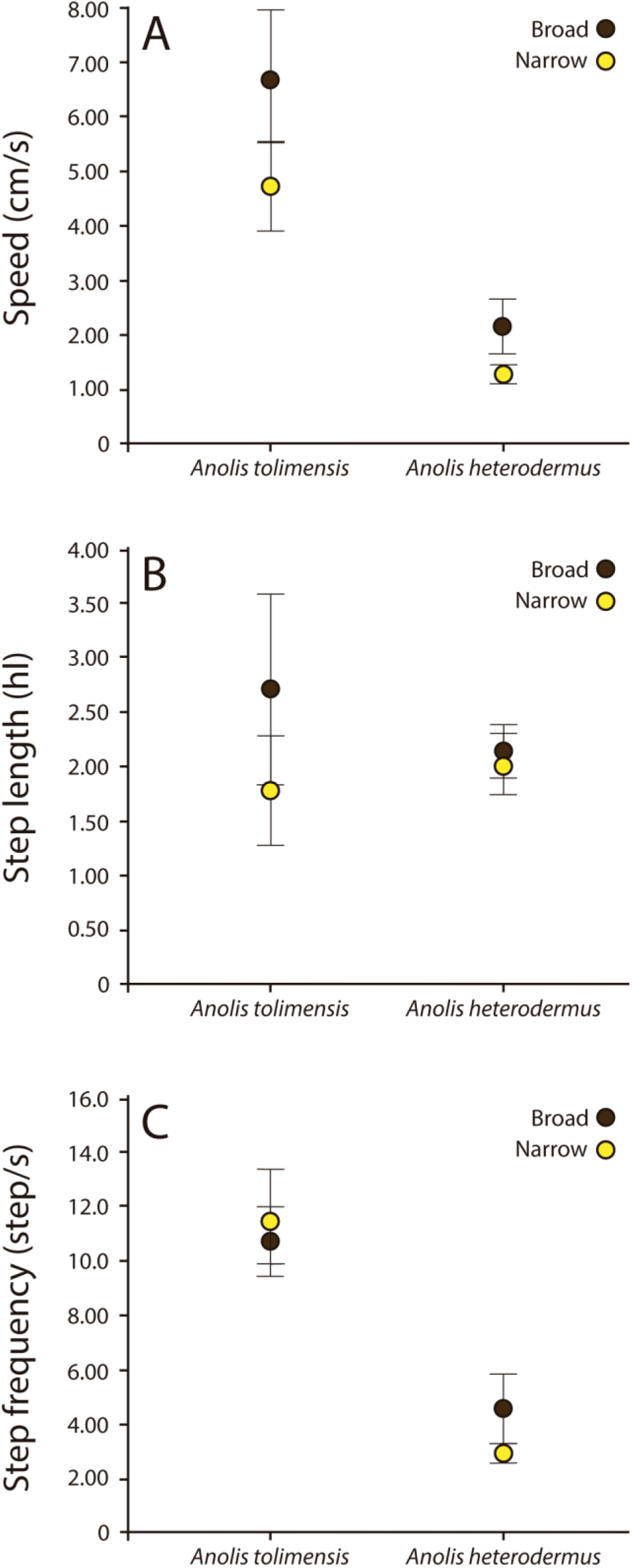
Values of speed, step length and step frequency obtained from *Anolis heterodermus* and *Anolis tolimensis* in wide and narrow substrates. The differences in the spatiotemporal characteristics of the movement performed by *A. heterodermus* and *A. tolimensis* in different substrates are shown. The data represent the mean and the standard deviation and correspond to the speed (**A**), the step length (**B**) and the step frequency (**C**) in both species.

We found that velocity and the step length are the most important variables that differentiate these species in their locomotor style, since *A. tolimensis* shows a greater step length for any substrate and it reaches speeds of up to 4.6 times higher than *A. heterodermus*. However, both species increase the step length and increase their average speed significantly on broader substrates (**Figure 6**).

## DISCUSSION

### Morphometric characteristics

In the general form of the body, differences can be found between the two species that were analyzed. *A. heterodermus* has a broad and robust head, short limbs with respect to its SVL and a compressed body, while *A. tolimensis* has a smaller head, long limbs with respect to its SVL and a cylindrical body. Both species differ in most of their body dimensions, except in the length of the humerus and radio-ulna. This could be related to the fact that *A. heterodermus*, which uses narrow surfaces, has a compressed and robust body like other arboreal lizards (Molnar et al. 2017), while *A. tolimensis* with a cylindrical body could be more stable on wider surfaces (Losos & Irschick, 1996; Zaaf et al. 1999; Herrel et al. 2008; Abdala et al. 2009).

These proportions coincide to the morphotypes assigned by Moreno-Arias & Calderón-Espinosa, (2016): the relative proportion of the hindlimb corresponds to less than 50% (approximately 35% in *A. heterodermus*) of its SVL, which is the morphotype MT4; and the relative proportion of the hindlimb is more than 80% (approximately 83% in *A. tolimensis*) of its SVL so it corresponds to the MT2 morphotype. Thus, the lizards located in the MT4 morphotype are those that exhibit the most extreme morphology, with short legs and arms associated with the use of very narrow surfaces and slow movements (Moreno-Arias & Calderón-Espinosa, 2016).

However, although the length and proportions of the limbs have been previously associated with habitat use and type of perch (Losos, 1990b, 1992), other morphological characteristics of *A. heterodermus* may be the product of ecological factors of the ecosystems where it has evolved. This species is the only *Anolis* in Colombia that is distributed at more than 3200 meters above sea level in the páramo (Miyata, 1983; Vargas-Ramírez & Moreno-Arias, 2014), which experiences drastic changes in temperature during the day, as well as high solar radiation and high water stress during the dry season (Wollenberg et al. 2014). The highland species generally has larger bodies and larger scales on the body to better withstand water stress (Soulé & Kerfoot, 1966), both characteristics present in *A. heterodermus*; and this explains the size differences with *A. tolimensis* that lives in low altitudes in Colombia.

### Appendicular Skeleton

The two species analyzed differ anatomically, both in the girdles and in the limbs. In the complex pectoral-sternum girdle *A. heterodermus* has a clavicle that is noticeably wider than in *A. tolimensis*, which increases the bone surface for greater muscle fiber insertion of the *Mm. clavodeltoideus pars superficialis* and *pars profundus*; these originate in the clavicle and interclavicle and are inserted in the anterior part of the humerus, promoting a greater force for the movement of the arm upwards and to the front (protraction and humeral abduction according to Herrel et al. (2008)) that *A. heterodermus* must develop in the narrow branches where it moves.

*A. heterodermus* has a concave coracoid at the base that is markedly convex anteriorly, whereas in *A. tolimensis* it is slightly convex at the base. In this bone the muscles of the *Mm coracobrachialis longus* and *brevis*, *M. coracohumeralis posterior* and *M. supracoracoideus* all originate; these are inserted into the humerus and allow the arms to approach the body (retraction and humeral adduction according to Herrel et al. (2008)). These differences of the coracoides are related to the greater mass of these muscles for *A. heterodermus* compared to *A. tolimensis*, allowing greater traction in climbing and in vertical movements. Also, in the coracoid region the biceps muscles (M. biceps I) are located. These muscles possess long and strong tendons, which in *A. heterodermus* are involved in keeping the body close to the grip surface to avoid possible falls, for example, due to wind and rain.

*Anolis heterodermus* has an elongated sternum that is consistent with the morphology reported by Herrel et al. (2008) for *A. valencienni*. The sternum of *A. tolimensis* is broad and short, as described in *A. sagrei*. The meso-sternum of *A. heterodermus* is long and thin and articulates only with two pairs of ribs instead of three as in *A. tolimensis*. These modifications are reported by Beuttell & Losos (1999) for other *Anolis* in the Carribean islands with climbing habits; and this characteristic could be related to a broad bone surface for the insertion of the humeral retractor muscles, such as *M. pectoralis pars superficialis*, which originates along the entire sternal plate, acting to resist gravity when climbing on vertical surfaces.

In the pelvic girdle we see a particular pattern between the continental and island species so far reported: *A. heterodermus* has a pelvic girdle with very broad elements, compared to *A. sagrei* and *A. valencienni* that exhibit relatively widened girdles (Herrel et al. 2008); and *A. tolimensis* has thinner bones than the other three species. Therefore, the species analyzed in this work differ from their ecomorphological equivalents in the Caribbean islands in the pelvic girdle, probably related to phylogenetic factors, but a comparative analysis with a greater number of species is required.

The widening of the pelvic girdle suggests a greater surface for muscle insertion related to the capacity of moving forward. For example, muscles associated with the pubis and ischium can lead to a greater pushing force in *A. heterodermus* compared to *A. tolimensis*, since they exhibit a greater mass (e.g. femoral adductor: *M. pubofemoralis pars ventralis*, *M. anterior ischiofemoralis*, *M. adductor femoris*, and femoral protractors: *M. pubofemoralis pars dorsalis int.*, and *ext.*, *M. ischiofemoralis dorsalis anterior*).

The knee extensors that originate in the ilium (e.g. *M. iliofibularis* and *M. A. pars dorsalis*, among others), could be involved on a horizontal surface in pushing the body forward, while on a vertical surface they would participate, in addition to the push, to overcoming gravity while climbing (James et al. 2007; Herrel et al. 2008). Thus, in *A. heterodermus* a wide pelvic girdle does not imply the use of muscles for more efficient movement on the ground. In contrast, in *A. tolimensis* a long ilium and a narrow pelvic girdle half laterally compressed, allows each leg to move easier in a sagittal plane (Russell & Bauer, 2008), and this represents an advantage for running.

The magnitude of the changes in *A. heterodermus* could be due to phylogenetic factors, since the *Phenacosaurus* clade has traditionally been associated with structural changes at the level of bones (Dunn, 1944; Lazell, 1969); whereas the absence of these characters in *A. tolimensis* could be due to the evolutionary history of their independent lineage.

*Anolis heterodermus* exhibits limbs with robust bones when compared to *A. tolimensis*. This contrasts with *A. valencienni* and other arboreal lizards that have thin and gracile bones, associated with the loss of body mass to counteract weight on inclined surfaces (Beuttell & Losos, 1999; Mattingly & Jayne, 2004, 2005; Herrel et al. 2008). It has been shown that the relative length and thickness of elements of the appendicular skeleton are associated with differences in locomotor abilities (Lemelin & Schmitt, 1998; Kirk et al. 2008; Patel, 2010; Almécija et al. 2015). The differences in *A. heterodermus* with respect to *A. tolimensis* could be due to a musculoskeletal specialization of the limbs, which allows a better performance of the grip (Zaaf et al. 1999; Moro & Abdala, 2004).

Other differences in the appendicular skeleton are observed in the autopodium. *A. heterodermus* has carpal elements and proximal epiphyses of metacarpals that are wider than *A. tolimensis*. A greater development of the proximal epiphyses of the metacarpals as a muscle insertion area is also a characteristic that is observed in arboreal species such as *Polychrus acutirostris* (Moro & Abdala, 2004) and *Chamaeleo calyptratus* (Molnar et al. 2017).

The absence of the first carpal bone in *A. heterodermus* was also seen, and although this feature has not been previously reported in other *Anolis* lizards, the reduction and modification of bone elements in the manus (within which there is a reduction of the palmar sesamoid and the central bone that are both characteristics present in both *A. heterodermus* and *A. tolimensis*) are linked to a more secure grip (Abdala et al. 2009; Fontanarrosa & Abdala, 2013; Molnar et al. 2017). According to Fontanarrosa & Abdala (2016), these characters are shared with other groups of lizards (*Iguana, Physignatus, Tropidurus, Anisolepis, Techadactylus, Gonatodes* and *Homonota*), and can be strongly linked to climbing and gripping skills. Thus, although *A. tolimensis* is not a strict climber, it has all the characteristics necessary for efficient climbing.

In the same way, the differences found in the tarsal elements, where the proximal tarsal bone of *A. heterodermus* has a very pronounced concave region with respect to *A. tolimensis*, could be related to a muscular and tendinous arrangement associated with the grip habit that requires greater muscular insertion surfaces in highly specialized lizards (Zaaf et al. 1999; Moro & Abdala, 2004; Abdala et al. 2009).

Other studies have found a strong phylogenetic signal associated with body characteristics such as the SVL and the proportion of the limbs (Moreno-Arias & Calderón-Espinosa, 2016; Poe & Anderson, 2019), and this could also have a phylogenetic incidence in the differences observed in the anatomy of the appendicular skeleton between *A. heterodermus* and *A. tolimensis*, associated with the evolutionary history of each lineage. This should be evaluated in a comparative analysis of the continental *Anolis*.

### Muscles and distribution of muscle mass

The results of this work show how the distribution of muscle mass varies between the two species in most functional groups. The two species differ considerably in terms of the total weight of the muscles in each limb. Although both species concentrate most of their muscle mass in the hindlimbs, *A. heterodermus* shows a 45% of total muscle mass in the forelimbs, while *A. tolimensis* only concentrates 30%. This distribution in both species is expected, since the hindlimb is typically dominant in the locomotion of lizards and is the one that generates the impulse for movement (Snyder, 1954; Reilly & Delancey, 1997a, 1997b; Irschick & Jayne, 1999).

However, it is clear that *A. heterodermus* distributes the muscle mass of the limbs almost equally, given a greater relevance of the forelimbs with respect to *A. tolimensis* and terrestrial lizards that distribute their weight mainly towards the hindlimbs (Herrel et al. 2008; Abdala et al. 2009). This may be due in part to the difference in body dimensions, since *A. heterodermus* has short hindlimbs, which reduce the total muscle surface.

Both species have the same total muscle mass with respect to their size, which indicates that the only difference is in the distribution of weight in the functional groups. Some functional groups show marked differences between both species, such as the retractor muscles of the femur, which are heavier in *A. tolimensis*. This could mean an increase in acceleration by having the ability to pick up the leg faster in a race event and with this would increase the frequency of steps in the movement phase (Mattingly & Jayne, 2004). This could also be seen in the performance tests, since *A. tolimensis* showed considerably higher speeds (4.6 times faster on average) than does *A. heterodermus*, where this functional group is less developed.

Likewise, *A. heterodermus* has heavier extensor muscles of the elbow and the abductors and adductors of the humerus when compared to *A. tolimensis*, and this could represent an advantage for supporting the weight of the body on vertical surfaces (Molnar et al. 2017). However, these modifications could reduce the possibility of moving quickly on both wide and narrow surfaces (Zaaf et al. 1999; Mattingly & Jayne, 2004, 2005).

When comparing the muscle mass of the functional groups of the species analyzed in the present work with the data previously published by Snyder (1954) for *Iguana iguana*, *Crotaphytus collaris* and *Sceloporus undulatus*, which are exclusively arboreal, terrestrial and semi-arboreal respectively, and Herrel et al. (2008) for island *Anolis*, we concluded that terrestrial lizards using broad substrates such as *A. tolimensis*, *A. sagrei* and *C. collaris* in general have similar muscle mass distributions. But they have certain variations in the femoral retractors, femoral adductors and ankle extensors (**Figure 7**), which may be associated with specific variations in the use of the microhabitat and its frequency of use. For example, *A. tolimensis* might use the soil less frequently than *A. sagrei*. The arboreal lizards show greater differences in most of the functional groups (**Figure 7**), accounting for the morphological variation at muscular level present in the species of this habit. Despite this, *A. heterodermus* was found to have more similarities with *A. valencienni* than with *I. iguana*, possibly due to its phylogenetic distance.

**Figure 7.**
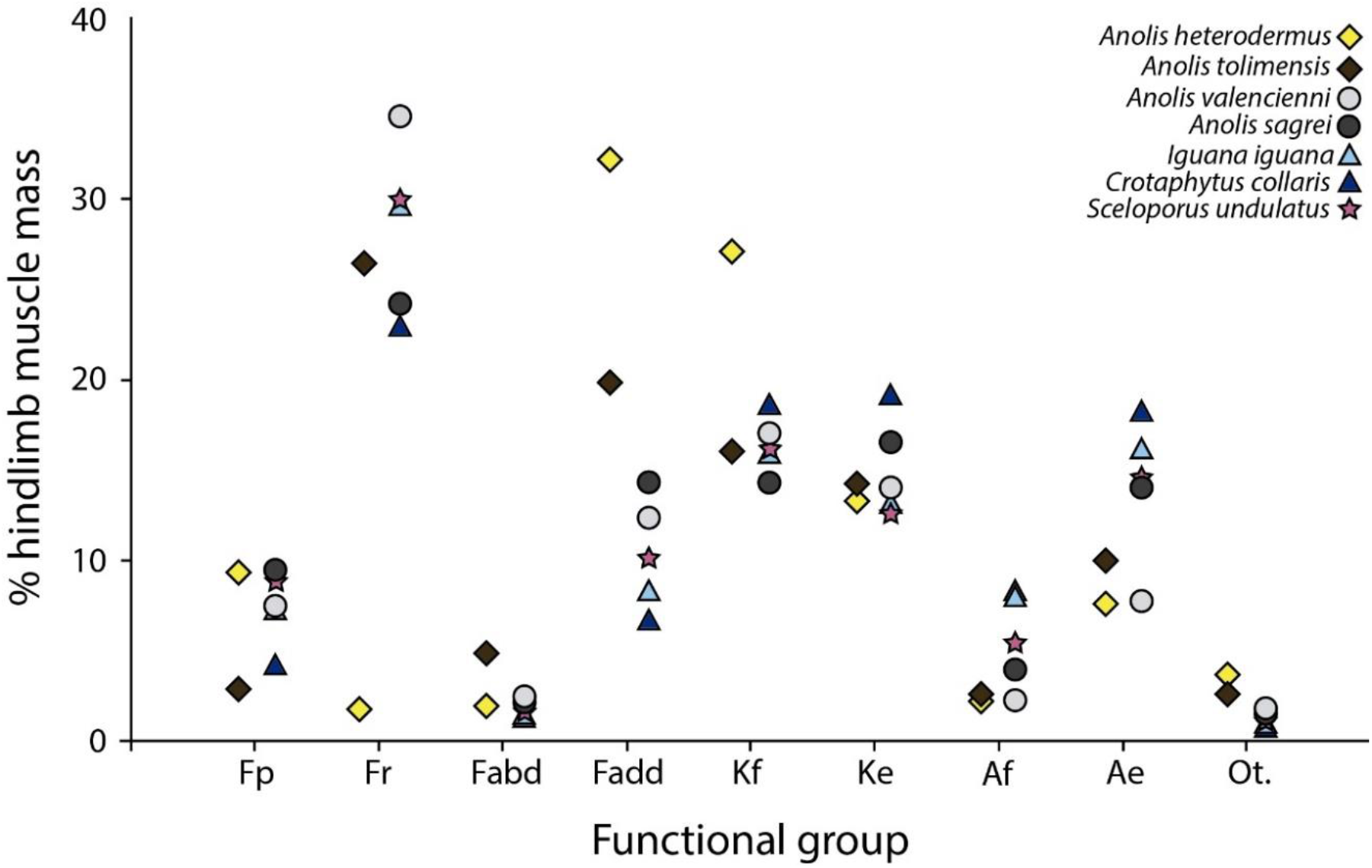
Distribution of muscle mass for some functional groups of the hindlimb in seven species of lizards with different habitat use. It shows the distribution of muscle mass of some functional groups in percentage. (**Fp:** femur protractors; **Fr:** femur retractors; **Fabd:** femur abductors; **Fadd:** femur adductors; **Kf:** knee flexors; **Ke:** knee extensors; **Af:** ankle flexors; **Ae:** ankle extensors; **Ot:** other hindlimb).

The myology and distribution of musculature in the Squamata are apparently very conservative (Diogo & Abdala, 2010; Molnar et al. 2017). *Anolis* and other closely related clades have no important differences in muscle distribution among lizards of terrestrial or arboreal habits (Lécuru, 1968; Russell, 1988; Zaaf et al. 1999; Moro & Abdala, 2004; Herrel et al. 2008), except in specific muscles such as carpal flexors, which are modified in some species in the insertion and with tendons that are more or less short between digits III and IV (Molnar et al. 2017; Lowie et al. 2018). In this case, both *A. heterodermus* and *A. tolimensis* do not exhibit significant differences in the appendicular muscle arrangement except for the muscles associated with the carpus in *A. heterodermus*, which have thicker and more conspicuous tendons than in *A. tolimensis*. This is especially true in the muscles *M. pronator accesorius* and *M. extensor digitorum longus pars profundus*, a feature not previously reported by Herrel et al. (2008) and Moro & Abdala (2004).

Similar to bone modifications in the carpus, these changes could be associated with specializations for gripping narrow surfaces (Fontanarrosa & Abdala, 2013; Molnar et al. 2017), as an additional modification in the tendinous pattern P of the palmar surface, where the tendon plate in the hand is reduced and the muscles of the digits derive from tendons accompanying the phalanges, a pattern found in both species. The P pattern has been associated with clades such as *Polychrus, Chamaeleo* and *Anolis* that grip to narrow surfaces (Moro & Abdala, 2004; Abdala et al. 2009). This would indicate that, although *A. heterodermus* is a specialist in grasping narrow branches, *A. tolimensis* also has the ability to move through this environment, since they exhibit the P configuration of the palmar surface. In a comparison between ground *Anolis*, *A. tolimensis* and *A. auratus* that have morphological and ecological similarities (although not in an equivalent way) (Moreno-Arias & Calderón-Espinosa, 2016), they share the same P configuration (personal obs), which could suggest that this arrangement associates both phylogenetic and functional factors, supporting the idea that *Anolis* are natural climbers.

In addition to this, as reported by Molnar et al. (2017) for chameleons, *A. heterodermus* has greater development of the humeral protractors with respect to *A. tolimensis*, which allows a more efficient movement of the shoulder in a parasagittal plane and favors the movement of the pectoral girdle in the direction of the body (Peterson, 1984) and, therefore, may imply an improvement in climbing. It is evident how *A. heterodermus* has tended to super-specialization of the forelimb, mainly of muscles and bones associated with the manus, which represents a significant advantage in the movement associated with the microhabitat that it occupies.

There are similarities in some functional groups used for common movements, such as the other forelimb and hindlimb muscles groups used for rotation and pronation among others, and the wrist flexors, which are apparently conserved in lizards (Lécuru, 1968; Russell, 1988; Zaaf et al. 1999; Moro & Abdala, 2004).

### Locomotor performance

The analyses show a relationship between the movement pattern of the two species with respect to the substrate, tending to have slower movements on the thin platform, similar to that found by Herrel et al. (2008). However, although both species move more slowly on thin surfaces, the data obtained in the analysis of *A. heterodermus* are considerably more homogeneous than those obtained from *A. tolimensis* (**Figure 6**). It is clear that *A. tolimensis* uses fast reaction movements with higher acceleration velocities, while *A. heterodermus* makes long and harmonic movements, moving at a much more constant rate. In similar tests applied to the arboreal lizard *Anolis equistris*, the results show that individuals do not change in the parameters of steps per second on both substrates (Abdala et al. 2009), and this could determine that the twig *Anolis* have a relatively low sprint speed, but its locomotion is safer (Losos & Sinervo, 1989; Losos & Irschick, 1996; Vanhooydonck et al. 2006b).

As verified by Herrel et al. (2008), the significantly lower reaction speed in *A. heterodermus* with respect to *A. tolimensis* can be attributed to the importance of maintaining the center of mass close to the surface at locomotion. When a lizard performs movements on wide surfaces, the center of gravity remains constant between steps due to the additional range of movement that exists laterally. However, when the movement is over narrow surfaces, having a higher reaction speed for explosive propulsion could be less optimal, since the lateral displacement induces a change of the center of mass away from the surface (Spezzano & Jayne, 2004). Thus, the more harmonic movements of *A. heterodermus* could mean an advantage that allows them to stay on narrow surfaces in a more efficient manner. In contrast, by more frequently using wider surfaces for foraging and breeding activities, *A. tolimensis* will necessarily depend on a high reaction rate to perform better on hot surfaces and as an antipredation strategy (Mattingly & Jayne, 2004).

Although the differences observed from the performance tests carried out between *A. heterodermus* and *A. tolimensis* agree with the results obtained by Herrel et al. (2008) in *A. sagrei* and *A. valencienni*, we must take into account the differential habitat use by *A. tolimensis*, which moves on wide surfaces most of the time and occasionally on narrow surfaces (Ardila-Marín et al. 2008; Moreno-Arias & Calderón-Espinosa, 2016). Due to the evolution of subdigital lamellae, *Anolis* lizards are arboreal in different degrees; and although they show great diversity in the use of microhabitat, many of them retain the ability to climb in specific situations (Losos, 2009).

The continental species differ ecologically from the Caribbean species (Irschick et al. 1997; Schaad & Poe, 2010). Although *Anolis* morphotype MT4 shows strong similarity with its respective ecomorph in the Caribbean, more than any other group (Schaad & Poe, 2010), it is not so with the MT2 morphotype, which is difficult to quantitatively compare with any of the insular ecomorphs given the ecological differences in the species (Moreno-Arias & Calderón-Espinosa, 2016). Thus, the species analyzed in this study do not exhibit completely contrasting life habits (e.g. *A. tolimensis* using branches occasionally) (Ardila-Marín et al. 2008), and this causes the differences observed in the locomotion patterns between *A. heterodermus* and *A. tolimensis* to be smaller in magnitude than those of *A. valencienni* and *A. sagrei* observed by Herrel et al. (2008).

## CONCLUSIONS

The results obtained in this article show that there are anatomical differences in the skeleton and the appendicular musculature of *Anolis* species that allow us to understand the locomotor mode of the species studied in relation to their use of the habitat. Although the species studied are phylogenetically separated, a morphological pattern is found that highlights the muscle specializations associated with habitat use in this genus.

It is evident that *A. heterodermus* exhibits specializations of the forelimb, associated with the muscles and bones of the manus (mainly of the carpus), which favors movement in the microhabitat that it occupies. Although *A. heterodermus* shows such specializations, many of the changes associated with the efficiency of vertical movement (climbing and grasping) were also observed in *A. tolimensis*, which shows how, although the *Anolis* species are associated with a specific microhabitat, they have the ability to move in arboreal habitats to a greater or lesser degree, supporting the idea that they are natural climbers.

The data provided by Herrel et al. (2008) and the present study, show how an understanding of the morphology of the locomotor apparatus can help to explain the evolutionary correlation of morphology, movement, locomotor style (advance characteristics) and habitat use in *Anolis* lizards. This demonstrating that morphological modifications based on habitat use go beyond simple external differences in the size and shape of the limbs.

## ACKNOWLEDGEMENTS

Thanks to Dr. Martha Calderón, curator of Reptiles Collection from the Instituto de Ciencias Naturales (ICN-UNAL), for the loan of the specimens, for her statistical advice and for her contributions in the revision of the manuscript. We thank Dr. Nelsy Pinto for the information on some specimens and Dr. Pedro Sánchez for his advice in the statistical processing of data. We also thank Laboratorio de Equipos Ópticos Compartidos of Departamento de Biología, Universidad Nacional de Colombia, Sede Bogotá, and Dr. Luis Carlos Montenegro for the loan of the necessary equipment for the elaboration of this work. We thank El Recodo farm and its administrative committee, the Villa Marcela Poultry Farm and the Castañeda Family, which allowed the captured specimens to be obtained. Finally, thanks to Miguel Méndez, Johana Muñoz and Miller Castañeda for their help in data collection and field work, Maria José Espejo for the anatomical English corrections and Dr. Thomas R. Defler for the English editing.

## AUTHOR CONTRIBUTIONS

All the authors worked equally in the elaboration of this manuscript.

